# *In situ* SAXS of protein deposits in Alzheimer’s disease

**DOI:** 10.1101/868273

**Authors:** Biel Roig Solvas, Bradley T. Hyman, Lee Makowski

**Affiliations:** Department of Electrical and Computer Engineering, Northeastern University; Massachusetts Alzheimer’s Disease Research Center, Massachusetts General Hospital; Department of Bioengineering, Northeastern University; Chemistry and Chemical Biology, Northeastern University

## Abstract

Deposits of Aβ peptides (plaques) and tau protein (neurofibrillary tangles (NFTs)) are ubiquitous features of brain tissue in Alzheimer’s disease. Their contribution to disease etiology remains controversial. The molecular-to-nano-scale organization of fibrillar species in these protein aggregates remains uncertain, but may contain clues as to the contributions of these structures to disease. Whether or not all plaques are the same structure, and all tangles are the same, has implications for current hypotheses about polymorphic templated misfolding of their constituent proteins, Aβ and tau. Here we use x-ray microdiffraction in the small-angle regime (SAXS) to probe the molecular organization of these deposits. Using unstained histological sections of human brain tissue, we demonstrate that SAXS can characterize Aβ fibrils and tau filaments *in situ*. Aβ fibrils have a cross-sectional radius of gyration (R_xc_) of ~45 Å, and larger (R_xc_ >150 Å) aggregates appear to represent Aβ fibrils that have coalesced side-to-side with one another to create fibrillar bundles or macrofibrillar aggregates. Tau fibrils exhibit an R_xc_ of ~55 Å with little sign of coalescence into larger structure. The *in situ* mapping of these structures revealed subtle variation in Aβ structure across different brain areas and different cases.

## Introduction

How deposits of Aβ peptides (plaques) and tau protein (neurofibrillary tangles (NFTs)) accumulate in Alzheimer’s disease (AD) is uncertain^1–3^. Senile plaques, composed largely of Aβ peptides, have been variously hypothesized to act as sources, sinks or reservoirs of toxic species^1,2^. Both lesions disrupt the brain areas in which they occur, although intra-neuronal neurofibrillary tangles (NFTs) comprised of tau protein appear more closely correlated with clinical status than Aβ plaque burden^4^. Dense, Aβ-containing neuritic plaques^5^ do disrupt the extracellular space where they are observed, associated with activated glia, synpase loss, and often are associated closely with intracellular tau pathology^4^ in dystrophic neurites, suggesting they also play an active role in neurotoxicity. A major goal of structural studies of these protein deposits is to relate their structural organization to their biology. One hypothesis suggests that these misfolded proteins are uniform across brain regions and across individuals, a consequence of a unitary fibrillar structure that has been described by ssNMR and, more recently, by cryo-EM. An alternative hypothesis is that the aggregates are structurally related but distinct enough from individual to individual (or even brain region to brain region) to support the idea of “strains” - that might have differing pathophysiological roles^6–8^. To begin to examine these issues, we used a conformation sensitive biophysical approach, scanning x-ray microdiffraction, which (unlike, for example ssNMR or cryo-EM) can report on structure in the spatial context of intact tissue, and without amplification of structures. Here, we show how scanning x-ray microdiffraction can provide information on the spatial location of Aβ and tau aggregates within unstained neural tissue and provide information on the molecular structures of these fibrils at much higher resolution than is possible with standard neuropathological methods.

The detailed structures of fibrillar assemblies of both Aβ and tau, either isolated from brain or *in vitro*-assembled, have been studied extensively, defining an ensemble of structures, some or all of which may be relevant to disease. All amyloid structures contain a core of prototypical cross-β structure in which the β-strands of the peptides are extended perpendicular to the fibril axis. Tau filaments have a cross-β core comprised of 72 of the 352-441 amino acids in the six tau isoforms, with the remaining bulk of the protein forming a largely unstructured fuzzy coat^9^. In Aβ amyloid the β-strands may fold into one of a number of different conformations within the protofilament. ssNMR studies of material extracted from human brain tissue and amplified *in vitro* suggest that any given fold of the peptide may be associated with a specific AD clinical subtype^7,8^. This has also been suggested by use of fluorescent ligands to probe the molecular structure of Aβ in plaques from patients who had died with distinct subtypes of AD^10^. However, the mechanisms by which specific structural forms of amyloid differ in the context of a particular clinical presentation remain unknown.

Previous studies have shown that polymorphism in structure of Aβ fibrils is observed at all length scales, including molecular-scale arrangement of peptides within individual protofilaments, the number of protofilaments making up a fibril, and their arrangement within the fibrils^11^. Whereas the peptide fold in a protofilament appears to be preserved during templated assembly of amyloid^8^, structural variations at larger length scales appear ubiquitous. This is apparent both *in vitro* and *in vivo*. Fibrils of Aβ peptides are formed by the side-to-side association of protofilaments that may twist around one another^7,11–14^. Variation in the pitch and breadth of *in vitro* assembled fibrils is sufficiently prevalent that it can be observed within individual electron micrograph fields of view^11^. *In situ*, x-ray microdiffraction has been used to characterize and map the spatial distribution of variations in fibril twist within and between plaques^15^. Fourier transform infrared micro-spectroscopy has been used to demonstrate conformational changes in Aβ structures prior to formation of amyloid plaques in transgenic mouse models of AD^16^. Whereas most amyloid fibrils appear to be 5-20 nm in diameter, depending on the number of constituent protofilaments^17^, much larger, tightly packed bundles of fibrils have been observed after long term incubation *in vitro*^18^ or in tomographic reconstructions of amyloid in a cell culture model of AD^19^. At still larger length scales, plaques exhibit a variety of morphologies. Although no standard nomenclature exists, most plaques can be classified as either diffuse, fibrillar or neuritic^5^, the latter being associated with a core of Aβ fibrils surrounded by dysmorphic, swollen neurites frequently containing aggregated tau proteins. The distribution of these distinct morphologies in neural tissue is frequently stereotypical^20^, and mapping the locations of polymorphic forms within the brain may provide clues as to the contributions that different structural forms make to disease progression.

The generation of molecular-level information on amyloid structure *in situ* is challenging. Three-dimensional reconstruction from cryo-electron micrographs requires isolation of the fibrils from brain tissues^9^. ssNMR requires isolation followed by *in vitro* templated assembly^8^. Both these processes destroy information relevant to the location and relative position of structures within the brain. Here, we take an approach that preserves information on spatial localization of the structures being studied. To do this, we use x-ray microdiffraction^15^ in the small-angle x-ray scattering (SAXS) regime for the mapping of amyloid polymorphisms *in situ*.

Scanning 18 μ thick histological sections with a 5 μ diameter x-ray beam, we probe the variation of fibril sizes and shapes within protein deposits embedded in human brain tissue. The x-ray scattering pattern from amyloid has a small angle (SAXS) and a wide-angle (WAXS) portion. Analysis of WAXS from micro-diffraction studies of AD^15^ provided information on the distribution of amyloid in unstained histological sections and provided insight into the polymorphism of structure at the level of the tilt and twist of protofilaments within and among plaques *in situ*. In this study, we focus on the SAXS portion of the scattering that contains information about the cross-sectional size and shape of fibrils and can map variations in these parameters as a function of position within the tissue.

## Results

### Mapping the Positions of Lesions in Unstained Tissue

Regions of tissue that exhibited strong immunostaining for tau or Aβ were selected for analysis by scanning x-ray microdiffraction carried out at the LiX beamline at the NSLS-II synchrotron at Brookhaven National Laboratory. The samples were mounted on sample holders compatible with the LiX beam line hardware and examined using an on-axis optical microscope mounted co-axial to the x-ray beam at LiX. Regions for detailed analysis using x-ray scattering of unstained sections were selected on the basis of the observed local burden of Aβ- or tau-stained lesions in serial sections. Regions of interest (ROI) were then examined with scans comprising 2500 (50×50) diffraction patterns, each probed with a 5 μ diameter x-ray beam, resulting in structural information over a 250×250 μ area. Observed intensity was considerably stronger in scattering from plaques than from surrounding tissue and the location and shape of plaques could be determined by a mapping of overall intensity. Samples containing only NFTs exhibited weaker scattering and the location of the NFTs could be determined equally well by a mapping of integrated intensities or apparent radius of gyration. NFTs exhibited a higher apparent radius of gyration than surrounding tissue. Figure 1 contains a mapping of apparent R_xc_ from a region of EC scanned by microdiffraction (left) and compares it to an optical micrograph of an immunostained serial section (right) that demonstrates the presence of tau-containing protein deposits in an immediately adjacent section.

**Figure 1.**
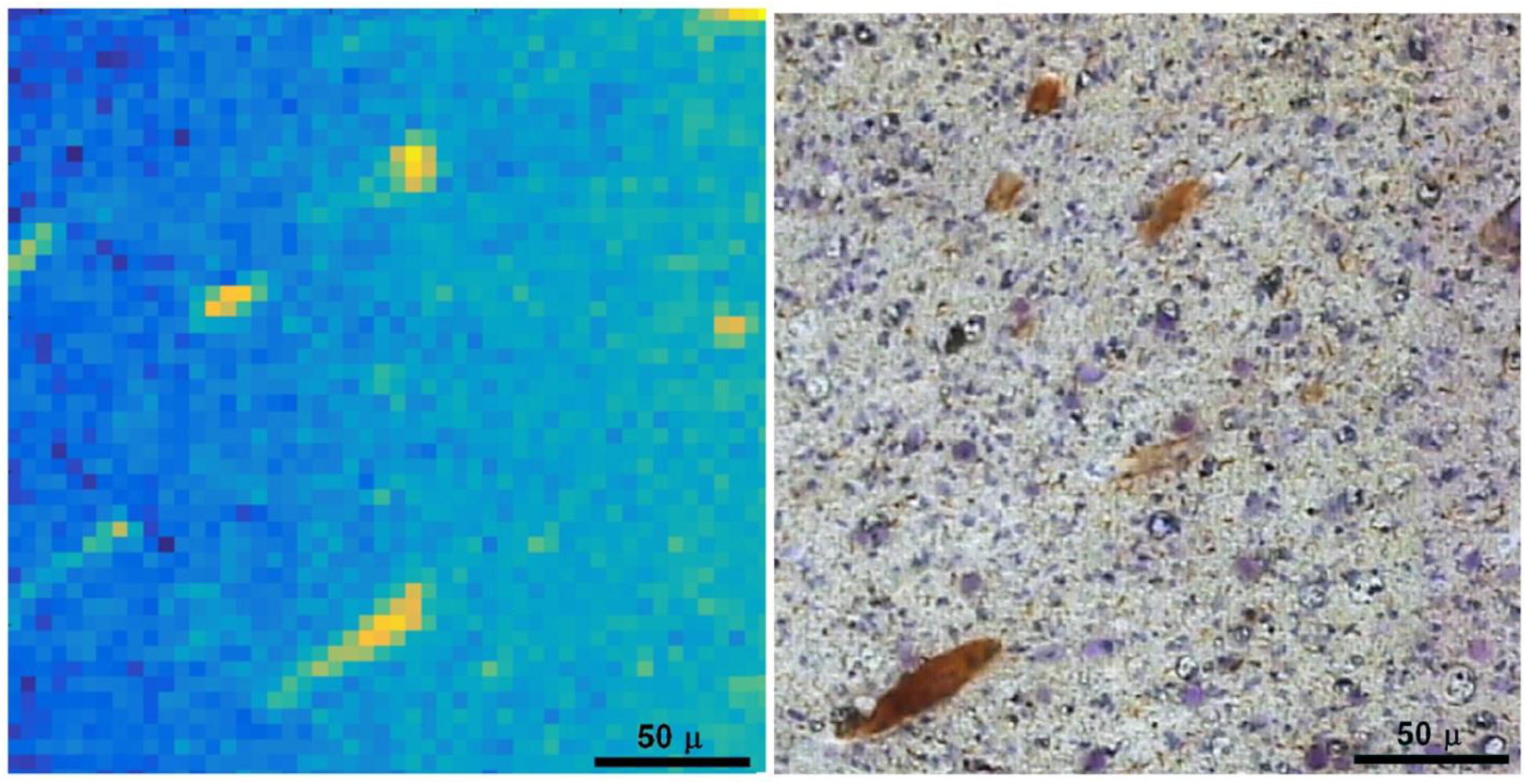
Detection of NFTs in an unstained tissue section of human entorhinal cortex (left) by mapping the apparent R_xc_ as a function of position in a scanned ROI (data from 2500 diffraction patterns). Comparison with immunostained serial section (right) from adjacent region indicated that regions rich in tau exhibit higher apparent radius of gyration than background tissue. Note that this is an adjacent section, so an exact 1:1 correspondence in the positions of NFTs in these images would not be expected. In the map of radius of gyration (left), yellow indicates apparent radius of gyration greater than that of background regions and corresponds to the location of NFTs

Figure 2 (top) shows contour plots of the integrated intensity of small-angle scattering in three scans, one each from EC, AC and FC. Figure 2 (bottom) are plots of apparent R_xc_ in the same regions, calculated from the circularly averaged data prior to background subtraction. This amounts to a mapping of the derivative of intensity at small angles and has proven to be a sensitive indicator of the locations of protein deposits. Small angle scattering intensities from lesions in the EC were essentially identical to one another, were considerably weaker than scattering from protein deposits in other tissues and were identified as NFTs by immunostaining. Scattered intensities from lesions in AC and FC exhibited greater variability in intensity profile, consistent with the presence of, variously, tau-containing filaments, Aβ fibrils, and larger aggregates.

**Figure 2.**
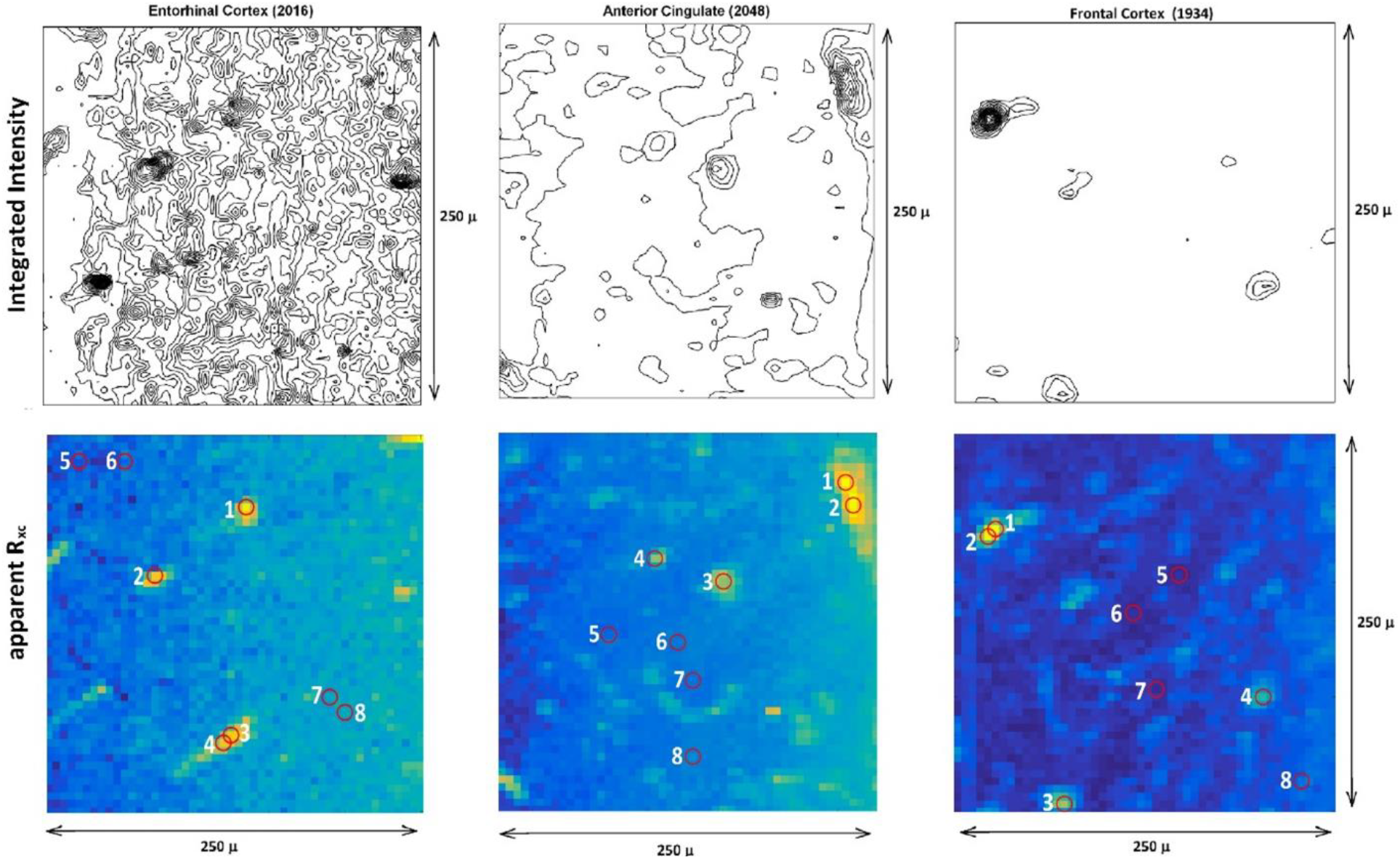
**(top)** shows contour plots of the integrated intensity of small-angle scattering in three scans, one each from EC, AC and FC. **(bottom)** are plots of apparent R_xc_ calculated from the circularly averaged data prior to background subtraction. Contour plots are drawn from data derived from 2500 scattering patterns collected on 50×50 grids with step size of 5 μ. Each field of view is 250×250 μ in size. The curves in Figures 3 and 4 are labeled to correspond to the locations of the lesions indicated in these maps. In cases where I(q) was virtually identical for different lesions, those data were averaged to improve the signal-to-noise ratio of the intensities. In data from EC and AC, positions 5-8 were considered background; for FC, positions 5-7 were considered background.

### Estimating the Size of Fibrillar Species

We observed that the SAXS data from most patterns could be fit to the sum of two exponential functions that represent Guinier approximations for scattering from fibrous materials with two different cross-sectional radii of gyration, R_xc_. In all cortical data sets studied, scattering could be interpreted as due to one population with an R_xc_ in the range of 35-60 Å and a second population with R_xc_ greater than 100 Å. All information available on the smaller structures indicates that the structures with R_xc_ ~ 45 Å are Aβ fibrils and those with R_xc_ ~ 55 Å are comprised of tau (see below). What about the larger structures?

Amyloid fibrils with lateral width > 250 Å have rarely been reported^11^, raising the question as to the identity of these larger structures. For a solid cylinder of radius, R, the R_xc_ = R/2^0.5^ meaning an R_xc_ ~ 100 Å corresponds to an outer radius of ~141 Å or lateral width greater than 280 Å. These large aggregates are the predominant structures in the tissue lesions in the FC (Table 1). They are distributed spatially in a way consistent with the expected spatial distribution of plaques in these tissues (Figure 2). Although larger than most amyloid fibrils that have been observed *in vitro*, the paucity of previous reports may be due to difficulties intrinsic to the isolation and detailed analysis of large bundles or coalesced aggregates of material. Furthermore, twisted bundles of fibrils have been assembled *in vitro*^18^ and bundles have been observed in a cell culture model of Alzheimer’s disease^19^. Since all aspects of the scattering data and neuropathology are consistent with their being composed of Aβ fibrils, our interpretation is that they represents aggregates of Aβ fibrils that have coalesced side-to-side to create fibrillar bundles or macrofibrillar aggregates.

**Table 1:**
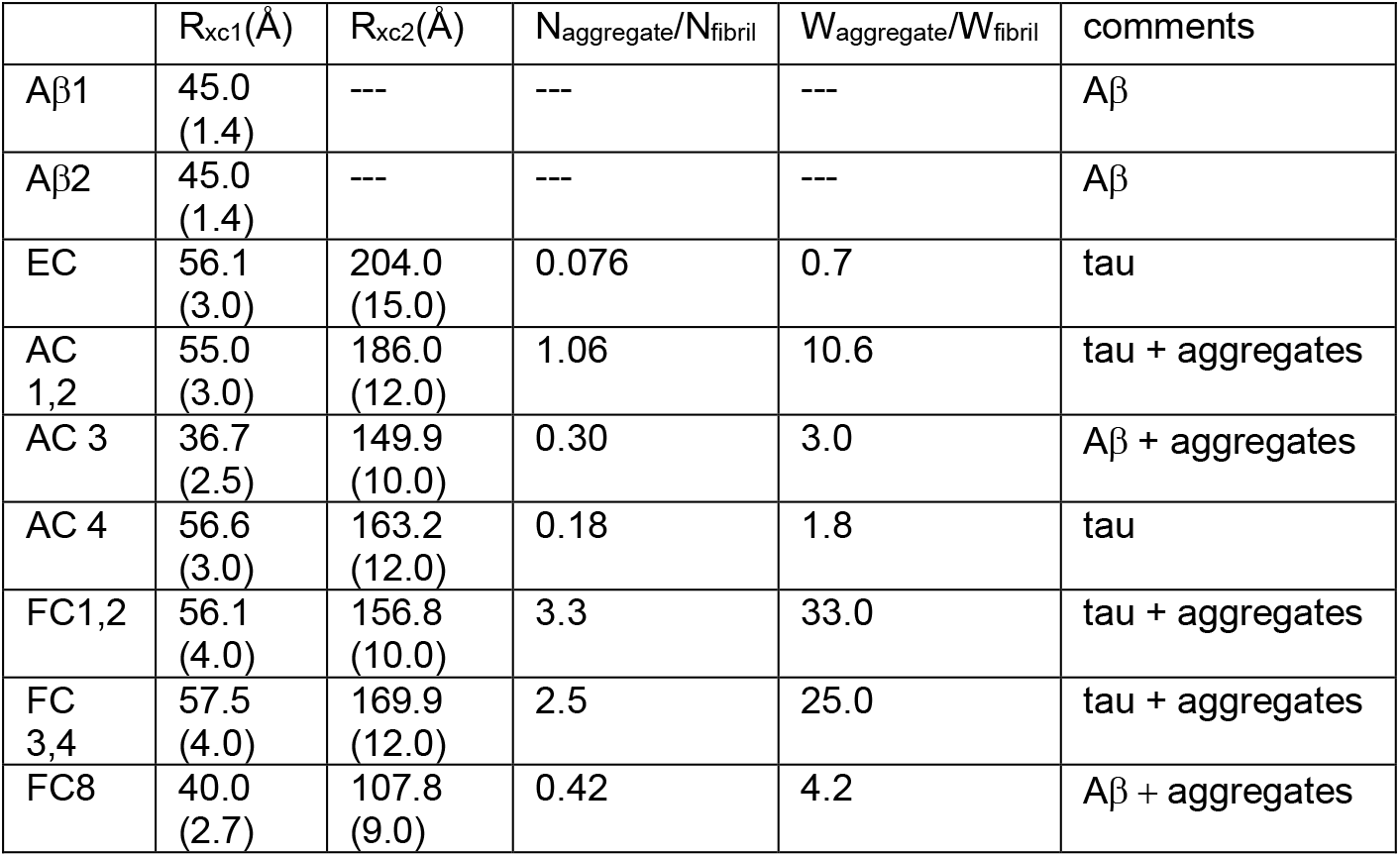
Data sets reported in this paper include those from *in vitro* assembled fibrillar preparations of Aβ40 peptides (Aβ1 and Aβ2); as well as data averaged from EC lesions (1-4); averaged from AC lesions 1 and 2; from AC lesion 3; from AC lesion 4; averaged from FC lesions 1 and 2; from FC lesions 3 and 4; and from FC lesion 8. Positions of each lesion in the ROIs scanned by microdiffraction are indicated in Figure 2. For each sample the data was fit to scattering from a small fibrillar structure and a large (R_xc_>100 Å) fibrillar structure. The R_xc_ of these two structures are indicated except for the *in vitro* assembled Aβ fibrils where no evidence of larger structures was observed. The relative intensities of the two Guinier approximations used to fit each data set was used to estimate the ratio of the number of fibrils to number of aggregates (N_aggregate_/N_fibril_) in each sample; and the mass ratio (W_aggregate_/W_fibril_) of the two structures in each sample/deposit. Numbers in parentheses are error estimates (see text).

Table 1 lists the R_xc_ for the patterns reported here. The scattering from *in vitro* assembled Aβ fibrils corresponds to an R_xc_ ~ 45 Å and no evidence for larger aggregates was present in those data. R_xc_ in the ~ 40 Å range was observed for two *in situ* deposits, one in the AC and one in the FC. We assign these lesions as containing Aβ amyloid fibrils. Scattering from all lesions analyzed in the EC corresponded to an R_xc_ of ~ 55 Å. Tau deposition in the EC has been observed to precede Aβ deposition^21^ and immuno-staining, as seen in Figure 1, indicated that this tissue contained only tau lesions and no Aβ amyloid, strongly suggesting that this scattering was due to fibrils composed of tau. Furthermore, the diameter of these fibrils is comparable to that observed by cryo-electron microscopy of tau filaments isolated from brain tissue^9^, supporting the assertion that they are tau. Similar R_xc_ were observed in scattering from some of the lesions in the AC and FC tissue, suggesting the presence of NFTs in those tissues. This is consistent with immunostaining that indicated these samples were positive for the presence of both Aβ and tau.

All data sets could be explained on the basis of the presence of a mixture of a homogeneous population of fibrils in the presence of larger aggregates. There is no reason to believe that aggregates within these lesions constitute a homogeneous population, but the goodness of fit to a sum of two exponentials as seen in Figure 3 suggests that the size distribution of the aggregates was limited. These larger aggregates are most likely formed by the side-to-side coalescence of smaller fibrils. The data, as observed, cannot be explained by the association of fibrils in random relative orientations as might be expected in the precipitation of fibrils from solution. Nor can it be explained by a loose bundling of fibrils. In each of those cases, the scattering fingerprint of the individual fibrils would dominate scatter with some modulation due to inter-fibrillar interference. It should also be noted that structures with R_xc_ > 500 Å would not be detected in these experiments due to the geometry of the x-ray camera used.

**Figure 3.**
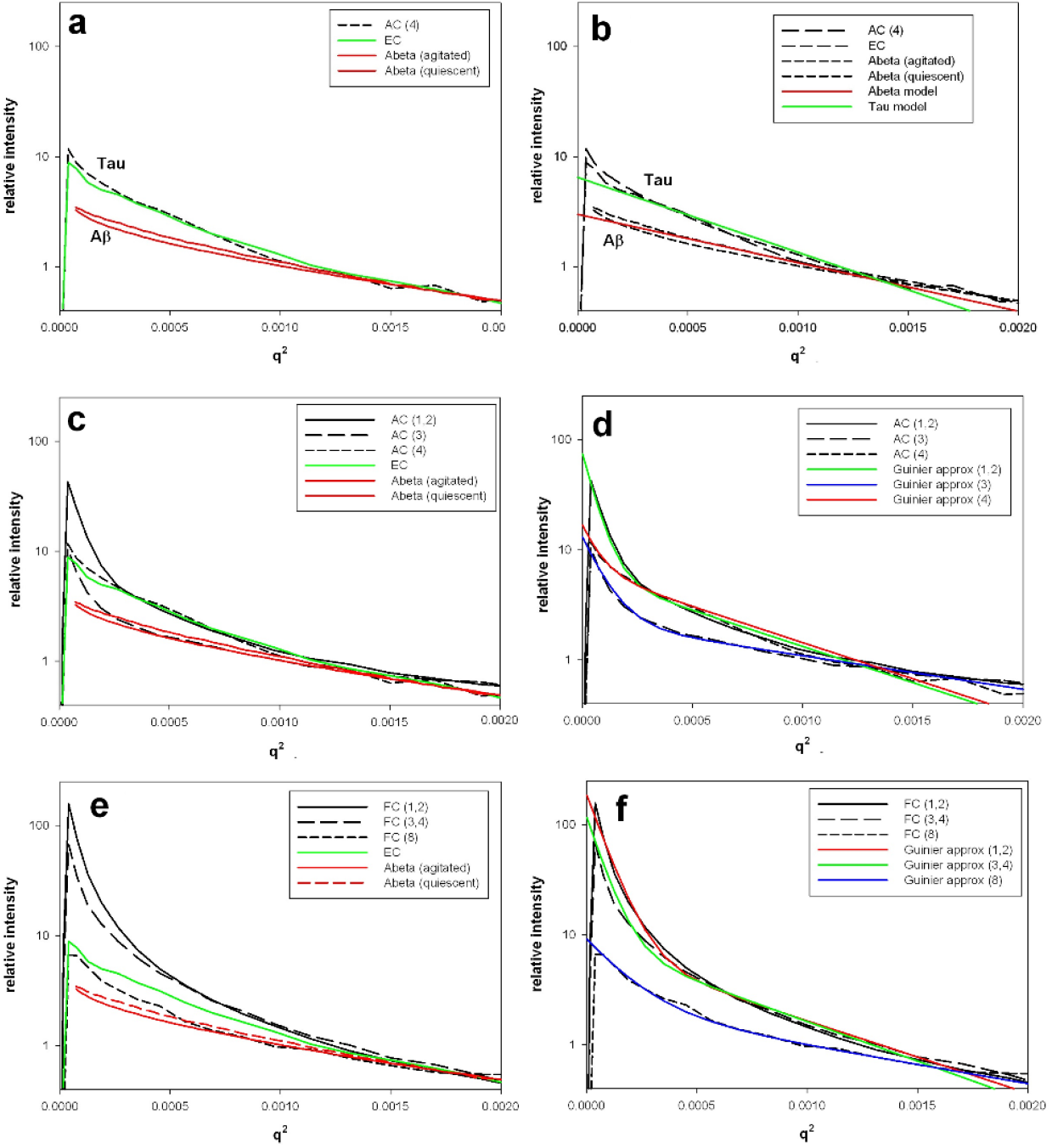
Guinier plots of intensity from *in vitro* and *in situ* fibrils: **(a)** Data from *in vitro* assembled Aβ fibrils (red) compared to data from *in situ* scattering from EC (green) lesions identified as tau by immunostaining and AC lesions exhibiting virtually identical scattering, suggesting that they also are comprised largely of tau. **(b)** Guinier approximations for the data in (a) demonstrating that the fibrils exhibit behavior as expected for a Guinier approximation of equatorial small-angle scattering with cross-sectional radius of gyration, R_xc_ approximately 55 Å for tau and 45 Å for Aβ. **(c)** Data from three lesions in the AC compared to the scattering from Aβ and tau filaments shown in (a). Data from lesion (4) in the AC suggest that tau is the dominant constituent. Data from the other fibrils indicate a mixture of fibrils with fibrillar aggregates as indicated by the presence of high intensity spikes at small scattering angles (low q). **(d)** Guinier approximations for the data in (c). Scattering from the lesions in AC (black solid and dashed lines) indicated a mixture of small fibrillar species and larger aggregates. Consequently, these data were fit (colored curves) by a sum of two Gaussian terms with parameters (I(q=0) and R_xc_) as listed in Table 1. **(e)** Data from three lesions in the FC compared to the data in (a). This comparison suggests that lesion (8) is largely comprised of Aβ fibrils, whereas the scattering from lesions (1/2) and (3/4) are difficult to assign since scattering from the aggregated material dominates scattering from individual fibrils. Best fit parameters derived from the Gaussian approximations as shown in Table 1 suggest that these are tau-based structures. Guinier approximations to the FC data are shown in **(f)**. All lesions in the FC appear to include large structures (R_xc_ ~ 150A), although these structures are less prevalent in lesion 8.

The ratio of the number of large fibrillar aggregates to the number of small fibrils can be estimated from the relative intensities of scatter from the small and large structures. This ratio varies widely over the lesions studied. In the FC, scattering from large aggregates dominates the observations, whereas in the EC, there is little evidence for the presence of large aggregates. The ***mass*** ratio of aggregates to individual fibrils is even more striking. The R_xc_ for the aggregates is roughly 3x that of the smaller fibrils. Since the mass of a fibril is proportional to 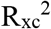, the large fibrils have of the order of ~ 10x the mass per unit length of the small fibrils, and, as detailed in Table 1 the mass of fibrillar aggregates in the two aggregate-rich regions of FC are ~ 25-35x that of the mass of individual fibrils in the same region. In other words, the vast majority of fibrils in these lesions is present as aggregated, macro-fibrillar structures.

Figure 3 provides a representative summary of the intensities (displayed as Guinier plots) observed. In the left panels of Figure 3, intensities observed from *in situ* measurements are compared to intensity from *in vitro* assembled Aβ peptides (red) and *in situ* scattering from tau aggregates (green) in the EC. In the right panels of Figure 3, Guinier approximations are displayed and compared to the *in situ* measurements.

### Distinguishing Tau from Aβ

Figure 3a compares the small angle scattering from one region of EC (demonstrated by immunostain to be NFTs) with scattering from a lesion in the AC and scattering from two samples of Aβ fibrils assembled *in vitro* under somewhat different conditions. The scattering from AC and EC samples are similar to one another but distinctly different from the Aβ scattering. In particular, as demonstrated in Figure 3b, the R_xc_ of individual fibrils in the AC and EC samples was approximately 55 Å; whereas the R_xc_ from the Aβ samples was about 45 Å. This strongly suggests that the NFT fibrils are distinctly different from the Aβ fibrils at the level of resolution attainable in SAXS. Furthermore, they indicate that protein deposits in the entorhinal cortex are largely made up of tau protein consistent with the results of immunostaining. They scatter weakly, are all very similar to one another, have an R_xc_ of ~ 55 Å and show little sign of coalescence into larger structures. This defines the most likely form of scattering from NFTs *in situ*.

### Anterior Cingulate

Figure 3c exhibits Guinier plots from AC and EC compared to scattering from *in vitro* assembled Aβ fibrils. The small-angle scattering from lesions in the AC exhibit three distinct patterns. In the first (- - -) scattering is indistinguishable from the lesions in the EC, strongly suggesting that the lesion is largely tau. In the second (- - -) the shape of the scattering curve suggests two structures – one indistinguishable from Aβ fibrils; the other indicating the presence of much larger structures (R_xc_ ~ 150 Å), most likely due to side-to-side aggregation of Aβ fibrils. The shape of scattering from the third lesion (------) again suggests the presence of two classes of structure – one about the size of the tau filaments; the other much larger with R_xc_ again > 150 Å. Interpretation of the scattering from this third lesion is not clear, but recalling that the size of the pixel is ~5 microns, a neuritic plaque may well be seen as having overlapping tau and Aβ signatures. If both NFTs and larger aggregates are present, the first thought would be that the larger structures are aggregates of tau. However, we see no sign of larger aggregates in samples containing only tau. Furthermore, neuritic Aβ plaques frequently adjoin degenerating axons and dendrites that contain aggregates of hyper-phosphorylated tau^4^ and may be involved in the induction of intracellular degenerative changes with tau pathology^21^. Consequently, a more likely interpretation of these data is that they arise from the co-localization of Aβ aggregates and NFTs. Even though Aβ is largely extracellular and tau intracellular, since the scattering volume is roughly 5×5×18 μ^3^, it is entirely possible that both structures be present within that volume. Estimates of R_xc_ in Table 1 are based on the comparisons shown in Figure 3d.

### Frontal Cortex

Immunostaining indicates the presence of both tau and Aβ in the FC tissue studied here. Scattering from these protein deposits exhibits significant diversity. Figure 3e displays plots of SAXS from this tissue. All lesions in the FC appear to include large structures (R_xc_ ~ 150 Å), although lesion 8 contains only a very small number of these aggregates (see Table 1). The scattering from lesion 8 suggests a mixture of Aβ fibrils with larger aggregates. The scattering from lesions 1/2 and 3/4 fit well to that expected from a mixture of NFTs and larger aggregates, similar to that seen in two of the lesions from the AC. Here, however, scattering from the larger structures dominates the observations, consistent with only a small fraction of the scattering being due to individual fibrils. Figure 3f provides the fit of the observed scattering to Guinier estimates of R_xc_ that are provided in Table 1.

### Cross-sectional Shapes

Reconstruction of the cross-sectional shapes of *in vitro*-assembled Aβ fibrils results in images that are consistent with cryo-electron microscope observations^22^ as shown in Figure 4. They indicate fibrils made of two protofilaments, each protofilament being composed of a pair of Aβ peptides every 4.7 Å along the fibril axis. This is consistent with the size and shape of *in vitro* assembled Aβ fibrils as observed in cryo electron microscopy^13,23^. As shown in Figure 4, reconstruction of cross-sectional shape from scattering of EC and AC lesions that exhibited an R_xc_ ~ 55 Å and minimal evidence of aggregates results in images of structures somewhat larger than the Aβ fibrils. Results of cryo-electron microscope studies of tau filaments isolated from human brain^9^ suggests that these R_xc_ ~ 55 Å structures are most likely comprised of tau. As explained in ‘Methods’, cross-sectional reconstruction from patterns containing scattering from both individual fibrils and aggregates is not possible.

**Figure 4:**
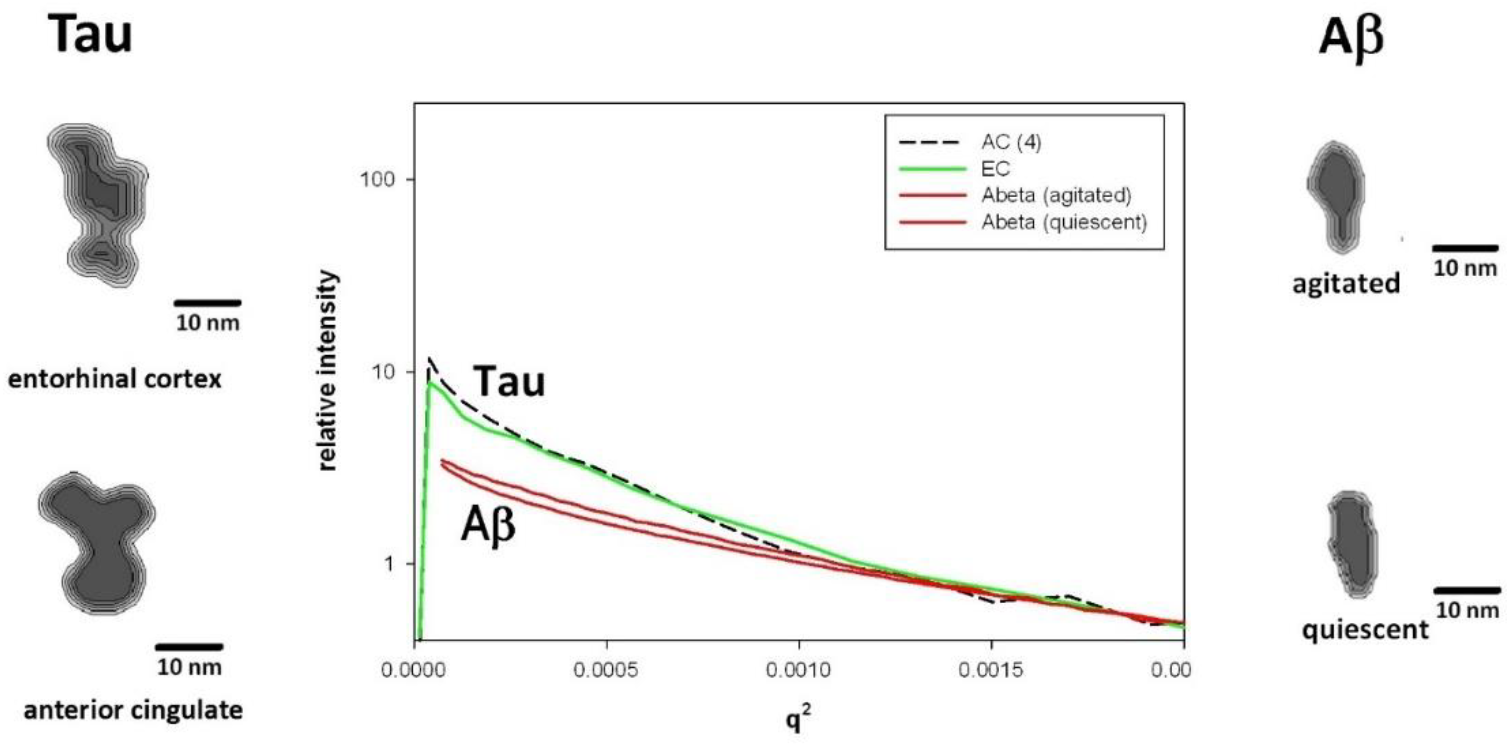
Reconstruction of fibril cross-sections from SAXS data collected from *in vitro*- assembled and *in situ* lesions (right and left, respectively) and the SAXS data used in their calculation (center). Two *in vitro* assembled Aβ preparations gave rise to slightly different scattering patterns (red curves). Comparison of reconstructions from these data with three-dimensional reconstructions from cryo-EM suggested that they were each made of two protofilaments but with the protofilaments arranged in slightly different ways. Scattering from lesions in the entorhinal cortex that stained for the presence of tau but not Aβ was interpreted as being due to NFTs (green curve). The close similarity of this scattering to that from some lesions in the anterior cingulate (dashed black line) strongly suggests that the AC lesions also were comprised of NFTs. Comparison with three-dimensional reconstructions from cryoEM data of fibrils isolated from human brain tissue suggests that the surface lobes of the reconstructed tau filaments (left) may correspond to the disordered ‘fuzzy coat’ (see Figure 5d of Fitzpatrick et al., 2017).

## Discussion

In this study we report the observation of x-ray scattering from protein deposits embedded within human brain tissue that preserves the spatial orientation and local microenvironment of the tissue, rather than requiring isolation, amplification, or the kind of restricted sampling and tissue preparation typical of electron microscopy. The protein deposits observed in AD brain exhibit characteristics consistent with those observed by electron microscopy from *in vitro-*assembled Aβ fibrils and tau filaments isolated from human tissue. The two classes of structures observed - those with R_xc_ ~ 45 Å and ~ 55 Å appear to correspond to fibrils composed of Aβ peptides and filaments of tau, respectively. This is the first time structural data has been obtained *in situ* on NFTs. The ability to directly observe these structures in unstained material, *in situ*, makes it possible to assess the variation of the molecular properties of these lesions across different neural tissues and across cases.

The observation of larger, R_xc_ ~ 150 Å, aggregates within protein deposits in these tissues indicates the presence of material that is composed of fibrils that are tightly packed, coalescing side-to-side to form large fibrillar aggregates or bundles that likely represents abundant constituents of neuritic plaques. This coalescence, or bundling of fibrils is consistent with the long-range correlation of fibril orientations reported to extend over >10 microns in dense plaques^15^ and the observation of bundles of Aβ fibrils *in vitro*^18^ and in a cell culture model of AD^19^. These observations suggest that very tight packing density is a common property of amyloid within dense cores of protein aggregates in human brain. In the tissues examined here we note that plaques in the neocortex are dominated by large aggregates of material and that these aggregates are less prevalent in the other regions studied. This might suggest that aggregate formation reflects the build-up of progressively larger, more tightly packed aggregates.

Preparation of histological thin sections preserves the anatomical topography, and allows individual lesions – rather than populations of lesions – to be identified and characterized.. Radiation damage from the microdiffraction has been observed in some cases as a bleaching of tissue observable by light microscopy. However, on re-scanning of a bleached region, the observed 4.7 Å scattering from the cross-β structures remains unchanged^15^. This is consistent with the interpretation that bleaching from x-ray exposure is due to damage to other tissue constituents and that the physical robustness of cross-β structures limits radiation damage to levels undetectable by small angle scattering.

The data presented here provide analysis of the classical lesions of Alzheimer’s disease in a size domain between cryo-EM and standard light histopathological methods, and have the advantage of being non-destructive and allowing interpretation of results in the context of the spatial orientation of intact tissues. We noted, for example, polymorphic Aβ structures, in accord with previous observations^7,8,10^. Moreover, we find that Aβ fibrils can coalesce into substantially larger structures to form a dense core plaque. This may give insight into the units that ultimately build and stabilize plaques.

SAXS studies also provided the new observations of NFT having a R_xc_ of ~55 Å in unstained tissue. While cryo-EM studies suggest that isolated fibrils from AD cases have similar structure^24^, that technique is, by necessity, highly selective. The current data are in accord with the idea that the fibrillar NFT structure is similar across regions and cases. Since SAXS is sensitive to multiple conformations of a protein in a heterogeneous mixture, our data reinforce the idea that the fibrillar forms of tau in NFT prefer a consistent and predominant molecular structure.

## Methods

Samples were prepared as previously described^15^. Briefly, human brain tissue was obtained from the tissue bank run by the Massachusetts Alzheimer’s Disease Research Center (MADRC) at the Massachusetts General Hospital. The brains collected by MADRC are handled, dissected, and stored in a uniform fashion. Brains are divided in half with one half fixed, processed in paraffin for neuropathological analysis and immunohistochemical studies and the other half generally snap-frozen in coronal slices. Age matched controls were between 85 and 95 years of age and showed no overt signs of dementia at time of death. Sections of fixed, paraffin-embedded tissue were cut 18 μ thick and captured on 12 μ thick mica sheets for x-ray microdiffraction experiments. They were subsequently de-paraffinized by heating at 75° C for a minimum of 12 hours and then washed for 5 minutes each with 100% xylene, 100% ethanol, 10% ethanol and water. Immediately before or after a section was cut for x-ray analysis, additional serial sections were cut to 5 μ thickness, captured on glass microscope slides, processed for histological examination and immunostained with 6F/3D antibody to locate amyloid plaques and DAKO A0024, a polyclonal total tau (non-phospho specific) antibody to locate NFTs. We collected SAXS data from regions of interest located in histological sections cut from samples that were seen by immunopathology to have heavy burden of plaque or NFTs. Data was collected from 12 regions of interest (ROIs) overall (30,000 diffraction patterns). Specific lesions were selected for detailed data analyses and those results are presented here. They are representative of the larger data sets collected throughout the study.

Samples of *in vitro* assembled Aβ_40_ protein were produced by methods detailed previously^8^ and kindly provided for these studies by Dr. Robert Tycko (NIH, Bethesda, Md.). Samples of fibrils formed by assembly under quiescent or agitated (stirred) conditions (as described^8^) were analyzed. X-ray microdiffraction in the SAXS regime was carried out at the LiX beam line (16-ID) at the NSLS-II synchrotron x-ray source at Brookhaven National Laboratory. The LiX beamline utilizes an undulator source with a Si(111) monochromator. The primary KB mirrors focus the beam to a secondary source and beryllium compound refractive lenses (CRLs) are used for secondary focusing. The x-ray energy range is 6-18 keV during normal operations. A beam size of 5 μ was used for the scanning measurements described here. A three detector configuration was used to collect both SAXS and WAXS data, the SAXS detector used with a 3000 mm sample-to-detector distance. Two tilted WAXS detectors were also used, but those data and their analysis are not described here. Data were collected on a 5 μ grid of scattering volumes with an exposure time of ~ 1 second. Scans that included 50×50 diffraction patterns (covering an ROI of 250×250 μ) required approximately 45 minutes collection time.

Scattering from regions of plaque exhibited greater intensity compared to adjacent tissue devoid of plaques. For each 2500-pattern scan, 10 or more patterns were identified as exhibiting scattering consistent with the presence of plaques or tangles; and 5-10 additional patterns were identified as exhibiting scattering suggestive of tissue absent of plaques or tangles. Intensities from all patterns chosen for detailed analysis were circularly averaged. Ten circularly averaged background patterns were averaged to generate a low noise estimate of background scatter and then subtracted from the patterns arising from plaques or tangles. The resultant background-subtracted intensities were multiplied by a geometric correction proportional to q (the momentum transfer - see below) to take account effect of disorientation of fibrils on the intensity of small-angle equatorial scattering^22^. Estimation of R_xc_ was made by fit of a sum of two exponentials to intensities over a q-range of 0.00625 to 0.05 Å^−1^. Errors were estimated by the range of parameters that resulted in fits within one standard deviation and are provided in Table 1 in parentheses as the largest deviation from the estimate provided that is consistent with the data.

Scattering from cross-β structures (including Aβ fibrils or tau filaments) is limited in most cases to equatorial intensities and intensities on layer lines spaced at ~ (1/4.7) Å^−1^. Consequently, any intensity in the small angle regime is solely due to equatorial scatter. The intensity of scattering on the equator, I_eq_(q) of a fiber diffraction pattern can be expressed as^22,25–27^.

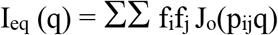

where f_i_ is the scattering factor of the ith atom, J_o_ is the zero-order Bessel function and q is the momentum transfer (q = 4πsin(θ)/λ; with 2θ being the scattering angle, λ being the x-ray wavelength) and p_ij_ the projection of the distance between atom i and atom j onto a plane perpendicular to the fiber axis. The sum is over all atoms in the structure. The form of this equation is analogous to the Debye formula for intensity in solution scattering, and it can be used to calculate a cross-sectional radius of gyration, R_xc_, and a cross-sectional shape through iterative refinement of two-dimensional shape analogous to the methods used to generate three-dimensional shape envelopes from solution scattering data^28–30^. Here we use methods as described previously^22^ to determine the R_xc_ and cross-sectional shapes of fibrils within the scattering volumes of tissue.

Although these methods worked well in samples containing homogeneous populations of fibrils, the approach broke down when applied to samples containing a heterogeneous population of fibrils. In many cases, the scattering we observed from lesions within human brain tissue was consistent with the presence of two distinct populations of structures within the scattering volume - one with cross-sectional diameter comparable to that expected for Aβ fibrils or tau filaments, and a second population of much larger structures. This motivated us to ask the question: If one tries to reconstruct cross-sectional shape using data arising from a mixture containing two distinct structures, what does the reconstruction look like? To answer this question, we calculated model scattering patterns expected for mixtures containing different ratios of cylinders with radii of 50 and 300 Å. Shapes reconstructed on the basis of model intensities calculated for homogeneous samples containing only small (or large) cylinders corresponded well to the cross-section of the cylinders used to generate the model intensities. However, when the same algorithm was used to reconstruct a cross-sectional shape from the sum of intensities from two cylinders of different radii with varying weight-ratios, the result was a highly irregular and variable shape that depended strongly on the starting point for the iterative reconstruction. Interestingly, the overall size of these irregular reconstructions corresponded to the size of the larger cylinder used to generate the model intensities; and the characteristic lengths of the irregular features in these cross-sections corresponded to the size of the smaller cylinder used to generate the model intensities. Although not representative of the cross-sectional shape of the cylinders used to compute the model scattering data, the dimensions of these reconstructions provide a manifestation of the dimensions of objects within the model scattering volume. We concluded, however, that these cross-sectional calculations were uninformative as to the shape of the larger structures and are consequently not reported here.

## Acknowledgements

We would like to thank Drs. Lin Yang and Shirish Chodankar for invaluable aid in collection of microdiffraction data at LiX and Dr. Robert Tycko for samples of *in vitro* assembled Aβ_40_ protein. The Massachusetts Alzheimer’s Disease Research Center is supported by NIH P30 AG062421. The LiX beamline is part of the Life Science Biomedical Technology Research resource, primarily supported by the National Institute of Health, National Institute of General Medical Sciences under Grant P41 GM111244, and by the Department of Energy Office of Biological and Environmental Research under Grant KP1605010, with additional support from NIH Grant S10 OD012331. As a National Synchrotron Light Source II facility resource at Brookhaven National Laboratory, work performed at Life Science and Biomedical Technology Research is supported in part by the US Department of Energy, Office of Basic Energy Sciences Program under Contract DE-SC0012704.

## Author Contributions

BRS contributed to selection of tissue; preparation of thin sections; collected SAXS data and carried out all preliminary data analysis. BTH provided overall project guidance and assistance in the use of the resources of the MADRC; LM had primary responsibility for all aspects of the project, helped in data collection; contributed to data analysis and wrote the manuscript.

